# Orthogonal analysis of mitochondrial function in Parkinson’s disease patients

**DOI:** 10.1101/2023.07.11.548533

**Authors:** Sander Barnhoorn, Chiara Milanese, Tracy Li, Lieke Dons, Mehrnaz Ghazvini, Stefania Farina, Daisy Sproviero, Cesar Payan Gomez, Pier G. Mastroberardino

## Abstract

The etiopathology of Parkinson’s disease has been associated with mitochondrial defects at genetic, laboratory, epidemiological, and clinical level. These converging lines of evidence suggest that mitochondrial defects are systemic and causative factors in the pathophysiology of PD, rather than being mere correlates. Understanding mitochondrial biology in PD at granular level is therefore crucial from both basic science and translational perspectives.

In a recent study, we investigated mitochondrial alterations in fibroblasts obtained from PD patients assessing mitochondrial function in relation to clinical measures. Our findings demonstrated that the magnitude of mitochondrial alterations parallels disease severity. In this study, we extend these investigations to blood cells and dopamine neurons derived from induced pluripotent stem cells reprogrammed from PD patients. To overcome the inherent metabolic heterogeneity of blood cells, we focused our analyses on metabolically homogeneous, accessible, and expandable erythroblasts. Our results confirm the presence of mitochondrial anomalies in erythroblasts and induced dopamine neurons. Consistent with our previous findings in fibroblasts, we observed that mitochondrial alterations are reversible, as evidenced by enhanced mitochondrial respiration when PD erythroblasts were cultured in a galactose medium that restricts glycolysis. This observation indicates that suppression of mitochondrial respiration may constitute a protective, adaptive response in PD pathogenesis. Notably, this effect was not observed in induced dopamine neurons, suggesting their distinct bioenergetic behavior.

In summary, we provide additional evidence for the involvement of mitochondria in the disease process by demonstrating mitochondrial abnormalities in additional cell types relevant to PD. These findings contribute to our understanding of PD pathophysiology and may have implications for the development of novel biomarkers and therapeutic strategies.

## Introduction

Parkinson’s disease (PD) is a high prevalence chronic neurodegenerative disorder. The disease is mostly idiopathic and its complex etiopathogenesis progresses along multiple, different, and typically converging biological pathways that include protein quality control, oxido-reductive homestasis, and intracellular trafficking (Betarbet, Canet-Aviles et al. 2006, Surmeier, Obeso et al. 2017). Within the intricate pathobiology of PD, anomalies in mitochondrial function are central and have been reported in both genetic and idiopathic PD cases, and there is consequently solid and general consensus on prominent role of mitochondrial dysfunction in PD etiopathogenesis (see e.g. (Bose and Beal 2016, Surmeier, Halliday et al. 2017, Mortiboys, Macdonald et al. 2018, Quansah, Peelaerts et al. 2018)). Reduced activity of respiratory complex I has been implicated in PD by evidence gathered at epidemiological level and in laboratory studies, as well as by clinical investigations that identified reduced peripheral complex I activity in patients’ platelets (Schapira, Cooper et al. 1989). More specifically, PD has been associated with the exposure to the complex I inhibitor rotenone in population studies (Tanner, Kamel et al. 2011), administration of this toxin in laboratory animals elicits a phenotype closely recapitulating that of the human disease (Betarbet, Sherer et al. 2000), and genetic deletion of a complex I essential subunit in mice causes a phenotype that closely recapitulates that of human PD (González-Rodríguez, Zampese et al. 2021). Moreover, several genes associated with familial forms of PD such as PARK2, PARK6, and PARK7 are involved in mitochondrial quality control (Beilina and Cookson 2016). These convergent data, obtained at both central and peripheral level, suggest that mitochondrial defects are systemic and causative of PD pathophysiology, rather than being a simple correlate. Such causative relationship constitutes the fundamental requirement for a valid surrogate endpoints (Fleming and DeMets 1996) and therefore indicates that mitochondrial biology holds potential both in terms of biomarker development and as targets for disease modifying therapies (Bose and Beal 2016, Macdonald, Barnes et al. 2018). Understanding mitochondrial biology in PD at granular level is therefore of primary relevance from both the basic science and translational standpoints.

To gather better insights on the relationship between mitochondrial function and PD, we recently analyzed mitochondrial function in fibroblasts from skin biopsies from a relatively large and well characterized cohort of idiopathic PD (iPD) patients (n=47) and correlated these biochemical parameters with both dopaminergic and non-dopaminergic clinical measures (Milanese, Payan-Gomez et al. 2019). To increase sensitivity toward mitochondrial defects, we used cell culturing conditions forcing energetic metabolism through oxidative phosphorylation (OXPHOS), which are not permissive for cells with severe mitochondrial defects and have been used as a diagnostic tool for mitochondrial diseases (Robinson, Petrova-Benedict et al. 1992). Using this simple, yet powerful stratagem we were able to show that alterations in mitochondrial function under these experimental setting correlate with clinical measures, and that the magnitude of mitochondrial alterations correlates with disease severity (Milanese, Payan-Gomez et al. 2019).

Few urgent standing questions, however, remain open. Evidence of peripheral mitochondrial dysfunction in PD, for instance, is discrepant and some studies failed to detect respiratory alterations in peripheral blood mononuclear cells (e.g. (Smith, Depp et al. 2018)). It is unclear whether this inconsistency is due a selective characteristic that limit mitochondrial defects to certain subsets/subtypes of PD, and/or if it rather depends upon the specific cell type that has been analyzed because of intrinsic cell dependent differences in bioenergetics. Correlation between mitochondrial respiration parameters and clinical measures is of potential translational relevance and should be validated on larger scale; it is however unknown whether correlation is detectable also in other peripheral cells than fibroblasts. Reconfirming the findings in cells different than fibroblasts would clarify these important aspects. Addressing these issue in blood cells would be highly desirable given they are easily accessible with non-invasive methodologies and would therefore greatly enhance the value of mitochondria in biomarker research. Bioenergetic studies in blood cells, however, pose critical issues because of the sharp differences in the metabolic layout of different cell type and even subtypes (O’Neill, Kishton et al. 2016). While purification via cell sorting methodologies may potentially circumvent this problem, it also comes with limitations regarding the amount of biological material available for the assays (Smith, Depp et al. 2018). A further fundamental question, which is essential to define in greater detail the role of mitochondrial dysfunction in PD, is whether peripheral mitochondrial anomalies reflect comparable alterations in neurons.

This study describes a parallel bioenergetic characterization of immature erythroid cells. i.e. erythroblasts, and iPSCs-derived dopaminergic neurons (iDAN) obtained from the very same and well characterized cohort analyzed in our previous study, and from the same patient. Moreover, it explores the correlation between clinical severity and mitochondrial alterations at transcription level in the large and well characterized PPMI cohort. The results reveal mitochondrial anomalies in erythroblasts and in iDAN. In peripheral cells, the observed reduction in mitochondrial respiration in PD peripheral cells is reversible because it can be potentiated in galactose culturing conditions. This effect, however, is not observed in iDAN, which therefore display a distinctive bioenergetic behavior. In summary, our study substantiates the relevance of mitochondrial biology in PD.

## Methods

### Human subjects

PD patients were recruited from the outpatient clinic for Movement Disorders of the Department of Neurology of the Leiden University Medical Center (Leiden, the Netherlands) and nearby university and regional hospitals. All participants fulfilled the U.K. Parkinson’s Disease Society Brain Bank criteria for idiopathic PD. The study was approved by the medical ethics committee of the Leiden University Medical Center, and written informed consent was obtained from all PD patients.

Fibroblasts were isolated at Leiden University Medical Center from skin biopsies derived from the ventral side of the upper leg and cultured under highly standardized conditions as previously described in (Milanese, Payan-Gomez et al. 2019). Peripheral whole blood was collected from PD patient’s at Leiden University medical Center and PBMCs isolated at the Department of Molecular Genetics at the Erasmus Medical Center in Rotterdam. Control iPSC were obtained from the Eramsus MC iPS Core facility.

Peripheral whole blood from 24 age-matched healthy controls (age >55 years) was obtained from Sanquin Rotterdam (NVT0585.00 Mantel, NVT0585.01 Annex).

Bioinformatic analysis was performed using the Parkinson’s Progression Markers Initiative (PPMI) database.

### Cell culture

To generate erythroblasts, PD patients’ peripheral blood mononuclear cells (PBMC) were extracted from 10 ml of freshly extracted blood with the use of Lympholyte-H (Cedarline) and Leucosep polypropylene tubes (227290, Greiner) according to manufacturer’s indications. Briefly, blood was diluted in PBS at a 1:2 ratio and loaded on a 15 mL Lympholyte – Leucosep tube. Blood was centrifuged at 800g for 25 minutes with no brakes at 4C. Upon removal of the plasma, the PBMC enriched cell fraction was collected, washed several times with sterile PBS and upon PBMCs were cultured in StemSpan SFEM medium (Stemcell Technologies) containing 2 mM Ultraglutamine (Lonza), 1% Nonessential aminoacids (NEAA), 1% penicillin/streptomycin, 50 ng/ml Stem Cell Factor, 2 U/ml Erythropoietin, 1 uM Dexamethasone (Sigma), 10 ng/ml Interleukin-3 (R&D Systems), 10 ng/ml Interleukin-6 (R&D Systems), 40 ng/ml IGF-1 (R&D Systems) and 50 ug/ml Ascorbic Acid (Merck) for 6-9 days refreshing half of the medium every other day starting from day 2. Erythroblasts were isolated when reaching 60-70% of the total cell population by gradient centrifugation at 1000 x g for 20 minutes at room temperature over Percoll (GE Healthcare). Isolated erythroblasts were frozen in FBS containing 10% DMSO at -80°C. Metabolic analysis was performed within 2 days after thawing.

### Generation of iPSCs

PD patients’ fibroblasts used in this study were prepared isolated at Leiden University Medical Center from skin biopsies derived from the ventral side of the upper leg and cultured under highly standardized conditions as previously described in (Milanese, Payan-Gomez et al. 2019). The study was approved by the medical ethics committee of the Leiden University Medical Center, and written informed consent was obtained from all PD patients.

Fibroblasts were reprogrammed to pluripotent stem cells using the CytoTune-iPS 2.0 Sendai Reprogramming Kit (A16517, Thermo Fisher) according to the manufacturer’s protocol.

### Generation of small molecule neural precursor cells (smNPCs)

Human iPSC lines were generated as previously described (Reinhardt, Glatza et al. 2013). Briefly, to generate embryoid bodies with neuroepithelial outgrowths (EBs), iPSC colonies were dissociated with 2 mg/mL collagenase IV and transferred to to non-adherent plates in hESC medum.

(Dulbecco’s modified Eagle’s medium (DMEM)/F12 (Thermo Fisher Scientific), 20% knockout serum (Thermo Fisher Scientific), 1% minimum essential medium/non-essential amino acid (NEAA, Sigma-Aldrich, St Louis, MO, USA), 7 nl ml−1 β-mercaptoethanol (Sigma-Aldrich), 1% L-glutamine (Thermo Fisher Scientific) and 1% penicillin/streptomycin (P/S, Thermo Fisher Scientific) supplemented with 10 μM SB-431542 (Ascent Scientific), 1 μM dorsomorphin (Tocris), 3 μM CHIR 99021 (Axon Medchem) and 0.5 μM Purmorphamine (Alexis) on a shaker in an incubator at 37 °C/5% CO2. On the second day, medium was replaced with N2B27 medium [DMEM-F12/neurobasal 50:50 (Thermo Fisher Scientific), 1% P/S, 1:100 B27 supplement lacking vitamin A (Thermo Fisher Scientific) and 1:200 N2 supplement (Thermo Fisher Scientific)] containing 10 μM SB-431542, 1 μM dorsomorphin, 3 μM CHIR99021 and 0,5 μM Purmorphamine. On day 4, N2B27 medium was replaced and supplemented with with 3 μM CHIR99021, 0,5 μM Purmorphamine and 150 μM Ascorbic Acid (Sigma).

On day 6, EBs were slightly triturated and plated on Matrigel-coated (Matrigel - 354277, Corning) plates at a density 10–15 EB per well containing smNPC expansion medium (N2B27 medium containing 3 μM CHIR 99021, 200 μM Ascorbic Acid, 0,5 μM Purmorphamine) and expanded for 5 passages before final differentiation. Medium was refreshed every other day.

### Differentiation of smNPC into midbrain dopaminergic neurons (iDANs)

smNPC were dissociated with Accutase at RT, diluted and seeded on Poly-D-lysine – Matrigel coated coverglasses in a 12 well plate at the concetration of 5×10^4^ cells per well in the Patterning medium [N2B27 medium containing 1ng/mL GDNF (Peprotech), 2ng/ml BDNF (Pepotech), 200 μM Ascorbic Acid and 0,5 μM Smoothened Agonist (SAG Pepotech)]. Medium was refreshed every 2 days.

At day 8, medium wash switched to the Maturation medium containing N2B27 medium, 2ng/ml GDNF, 2ng/ml BDNF, 1 ng/mL TGF-b3 (Peprotech), 200 μM ascorbic Acid and 5ng/ml of ActivinA for the first feeding and 2ng/ml ActivinA for the following feedings. Medium change occurred every third day.

### Flow cytometry

Erythroblasts and iPSCs were washed and resuspended in FC buffer (HBSS w/o calcium and magnesium + 0.5% BSA) and incubated with PE Mouse Anti-Human CD44 antibody (1:25, BD, 550989), FITC Mouse Anti-Human CD71 antibody (1:50, BD, 555536), or APC Mouse Anti-Human CD117 antibody (1:50, BD, 561118) for 30 minutes at 4°C. Mitochondria were stained with Mitotracker Green FM (100 nm, Cell Signaling, 9074) and active mitochondria with TMRM (100 nm, Thermo Fisher, T668) for 30 minutes at 37°C. Cells were detected by flow cytometry using a LSRFortessa Cell Analyzer (BD, USA). Flowjo software (BD, USA) was used for data analysis.

### Bioenergetics Assays

Oxygen consumption rates (OCR) and extracellular acidification rate (ECAR) were measured using a XF-24 Extracellular Flux Analyzer (Agilent Technologies), as previously described (Milanese, Payan-Gomez et al. 2019). Erythroblasts were seeded at a density of 2 × 105 cells/well on Cell-Tak (Corning, 354240) coated Seahorse plates in unbuffered XF DMEM medium (Agilent Technologies) supplemented with 1 mM sodium pyruvate, 2 mMglutamine and 10 mM glucose or galactose. Immediately after seeding, cells were centrifuged at 200 g for 1 minute to attach evenly to the bottom of the well and the plate was equilibrated for 30 minutes at 37°C in the absence of CO2. iPSCs derived from fibroblasts of PD patients and healthy controls were seeded at a density of 8 × 103 cells/well on Seahorse plates and differentiated to dopaminergic neurons over a period of 3 weeks according to the described methodology. On the experimental day, medium was changed to unbuffered XF DMEM medium supplemented with 1 mM sodium pyruvate, 2 mM glutamine and 10 mM glucose or galactose. Cells were incubated for 1 h at 37°C in the absence of CO2, before the Seahorse assay. For each assay, medium and reagent acidity were adjusted to pH 7.4 on the day of the assay, according to manufacturer’s procedure. Optimal cell densities were determined experimentally to ensure a proportional response to FCCP (oxidative phosphorylation uncoupler).

After 3 measurements to detect the oxygen consumption ratio baseline, cells were then challenged with sequential injections of mitochondrial toxins: 1 μM oligomycin (Adenosine triphosphate – ATP - synthase inhibitor), 1 μM FCCP, and 1 μM antimycin (complex III inhibitor). A minimum number of 5 replicates were performed for each cell line; data represent the mean of the different replicates. Basal respiration (measured as the average OCR rates at the baseline), maximal mitochondrial respiration (maximal respiration), reserve capacity (difference between maximal respiration and basal respiration) and respiration dedicated to ATP production (difference between basal respiration and oligomycin-dependent respiration) were used to investigate mitochondrial bioenergetics. Basal glycolysis, measured as extracellular acidification rate (ECAR) maximal glycolysis and reserve glycolytic capacity (difference between maximal glycolysis and basal glycolysis) were taken in account to investigate glycolytic properties.

### Immunofluorescence

Reprogrammed iPSCs cells cultured in a 8-chamber slide were fixed with 4% PFA for 15 minutes at room temperature. After incubation in ice cold methanol for 10 minutes cells were permeabilized in 0.1% Triton in PBS for 10 minutes and blocked using 1% BSA in PBS/0.05% Tween-20 for 30 minutes. Next, cells were incubated with primary antibodies diluted in blocking buffer overnight at 4°C - Mouse Anti-Human TRA1-81 (1:75, Abcam, AB16289#20), Rabbit Anti-Human OCT4 (1:250, Abcam, AB19857#8), Goat Anti-Human NCAM (1:100, R&D, AF2408), Goat-Anti Human SOX17 (1:100, R&D, AF1924) or Mouse Anti-Human Beta-Tubulin (1:1000, Merck, T8660) primary Chicken Anti-Human MAP2 (1:2000, Abcam, AB5392), Mouse Anti-Human TH (1:200, Millipore, MAB318). After washing with PBS cells were incubated with respective secondary Goat Anti-Mouse Alexa Fluor 546 (1:500, Invitrogen, A21045), Goat Anti-Rabbit Alexa Fluor 488 (1:500, Invitrogen, A11008#8a), Donkey Anti-Goat Alexa Fluor 488 (1:500, Invitrogen, A11055) or Goat Anti-Mouse Dylight 594 (1:500, Jackson, 115-515-166#7) antibodies diluted in blocking buffer for 1 hour at room temperature. Nuclei were stained with Hoechst 33342 (1:1000 in PBS, Thermo Fisher) for 10 minutes. Cells were next washed with PBS, mounted with ProLong Diamond Antifade Mountant (P36965, Thermo Fisher) and imaged using a Leica Stellaris5 confocal microscope.

### Bioinformatic Analysis of the PPMI dataset

Blood transcriptome data from the Parkinson Progressive Markers Initiative (PPMI) cohort (PPMI website: https://ida.loni.usc.edu/pages/access/geneticData.jsp#441) were obtained. The libraries were prepared using the NEB/Kapa (NEBKAP) based library prep, with second-strand synthesis. RNA sequencing was performed at Hudson Alpha’s Genomic Services Lab on an Illumina NovaSeq6000, generating 100 million 125 bp paired reads per sample. The Salmon files were imported into R using Tximport. To identify differentially expressed genes between PD groups and controls, the DESeq2 package was used. Normalized counts were subjected to Rlog transformation to improve distances/clustering for the principal component analysis (PCA). The cohort of subjects was divided into subgroups based on the delta-UPDRS-III (MDS-Unified Parkinson’s Disease Rating Scale, UPDRS-III at last visit - UPDRS-III at first visit) of PD subjects: those with a delta-UPDRS-III less than 0 (defined as mild) and those with a delta-UPDRS-III greater than 0 (defined as severe), as well as controls (CTRL). A threshold of significance at FDR < 0.05 was applied.

Gene Set Enrichment Analysis (GSEA) was conducted on an unfiltered, ranked list of genes. The analysis involved various terms from the Kyoto Encyclopedia of Genes and Genomes (KEGG), Reactome Pathway Databases, Hallmark Gene Set Collection, and WikiPathways (GSEA website: http://www.gsea-msigdb.org/gsea/msigdb/collections.jsp). Pathway information was obtained from the Kyoto Encyclopedia of Genes and Genomes (KEGG) available at the Molecular Signatures Database (http://www.broadinstitute.org/gsea/msigdb/index.jsp) or from the Hallmark Gene Set Collection (http://www.gsea-msigdb.org/gsea/msigdb/collections.jsp). Gene set enrichment with FDR < 0.1 was considered significant. Genes in each PD group were ranked based on the level of differential expression using a signal-to-noise metric and a weighted enrichment statistic.

Transcriptomic analysis was performed using R Studio version 4.2.3. The experiments were conducted with a minimum of three independent biological replicates. GraphPad Prism version 9 (GraphPad Software, La Jolla California USA) was used for all statistical analyses and graphical representations. P values were denoted as *P < 0.05, **P < 0.01, ***P < 0.001 and were considered significant. In the absence of indications, comparisons should be considered non-significant. Comparisons between two groups were analyzed using unpaired two-tailed Student’s t-tests, and comparisons between more than two groups were analyzed using either one-way or two-way ANOVA followed by Bonferroni multiple comparison post-hoc test.

## Results

### Mitochondiral bioinformatic analysis

We took advantage of the PPMI cohort to study expression of mitochondrial genes in peripheral mononuclear blood cells (PBMCs) from 393 iPD patients and healthy controls examined at two different time points, i.e. at the intake visit (i.e. visit 1), shortly after diagnosis, and in a follow up visit, after 36 months (i.e. visit 8). (Suppl.Table 1). Of the 340 of the 393 iPD patients and 159 of the 189 healthy controls that were part of visit 1 were also examined at visit 8. Given the intrinsic variability in PD progression, we divided the 340 patients examined at both visit 1 and 8 in two groups based on motor symptoms progression (i.e. UPDRS part III): mild cases included patients who did not show significant deterioration of motor symptoms (i.e. deltaUPDRS-III=UPDRS-III^visit8^-UPDRS-III^visit1^ ≤0), severe cases, encompassed patients with aggravation of motor symptoms who (i.e. deltaUPDRS-III=UPDRS-III^visit8^-UPDRS-III^visit1^ >0) (fig.1A).

**Figure 1.**
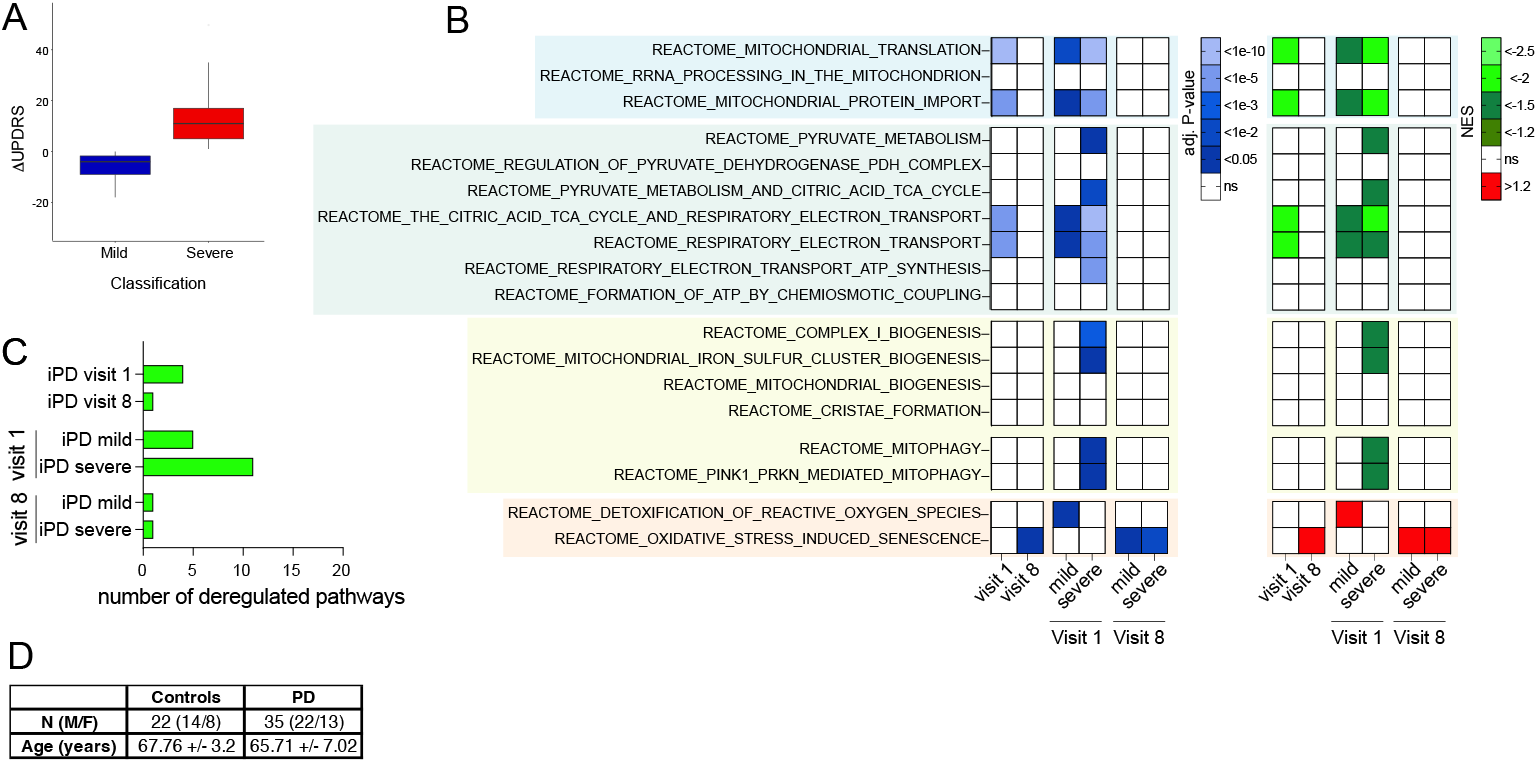
Gene set enrichment analysis reveals mitochondrial alterations in PD patients that parallel disease progression severity. (**A**) iPD in the PPMI cohort were stratified based on the evolution of the UPDRS-III score between visit 1 and visit 8. (**B**) GSEA indicates that multiple mitochondrial pathways are downregulated in iPD, at visit 1. Deregulation of mitochondrial pathways was not detected at visit 8, when pathways related to reactive oxygen species were instead upregulated. (**C**) At visit 1, patients who will develop more severe symptoms display deregulation in a higher number of mitochondrial pathways.

Over-representation analysis (ORA) based on significantly differentially expressed genes (DEG) (Boyle, Weng et al. 2004) did not highlight any differences between iPD and control, indicating that the disease is not associated to a pronounced mitochondrial defect in blood cells. To unravel more subtle differences, we took advantage of gene set enrichment analysis (GSEA) (Subramanian, Tamayo et al. 2005), which rather than focusing of single significantly DEG calculates aggregated statistical significance in a set of related genes with common biological function, i.e. a pathway.

We first analyzed PD patients as a single group, i.e. without stratifying for disease progression rate. Here, GSEA detected downregulation of mitochondrial patways involved in bioenergetic processes and mitochondrial related macromolecular synthesis (fig.1B).

We next stratified patients based on rate of disease progression. At visit 1, patients who subsequently developed more severe motor signs (i.e. severe cases) displayed downregulation in more mitochondrial pathways than patients with more benign disease progression (i.e. mild cases) (fig.1B,C). Deregulation pathways included bioenergetic processes, mitophagy, mitochondrial biogenesis, and mitochondrial related macromolecular synthesis. Interestingly, at visit 8 no deregulation of mitochondrial pathways was detected and processes related to reactive oxyugen species were instead upregulated (fig.1B). Collectively, these elements obtained in a large longitudinal PD dataset indicate that mitochondrial alterations are detectable at earlier rather than latear stages of the PD symptomatic phase, and also infom on disease progression.

### Blood cells

Encouraged by the supportive bioinformatic evidence obtained in the PPMI cohort, we next sought to determine alteration in mitochondrial function in blood cells. Because of the different metabolic layout of blood cell types (O’Neill, Kishton et al. 2016), bioenergetic analyses in this tissue should be performed in homogeneous and characterized populations. We followed a novel strategy and investigated mitochondrial function in immature erythroid progenitors (erythroblasts) obtained from peripheral blood mononuclear cells (van den Akker, Satchwell et al. 2010). This approach does not require preliminary antibody-based cell sorting and allows easy expansion of erythroblasts, with consequent generation of large and very homogeneous population from limited amount of starting material (fig. 2C) (Leberbauer, Boulmé et al. 2005, van den Akker, Satchwell et al. 2010). Erythroblasts retain functional mitochondria, which will be lost during terminal differentiation (Liang, Menon et al. 2021).

**Figure 2.**
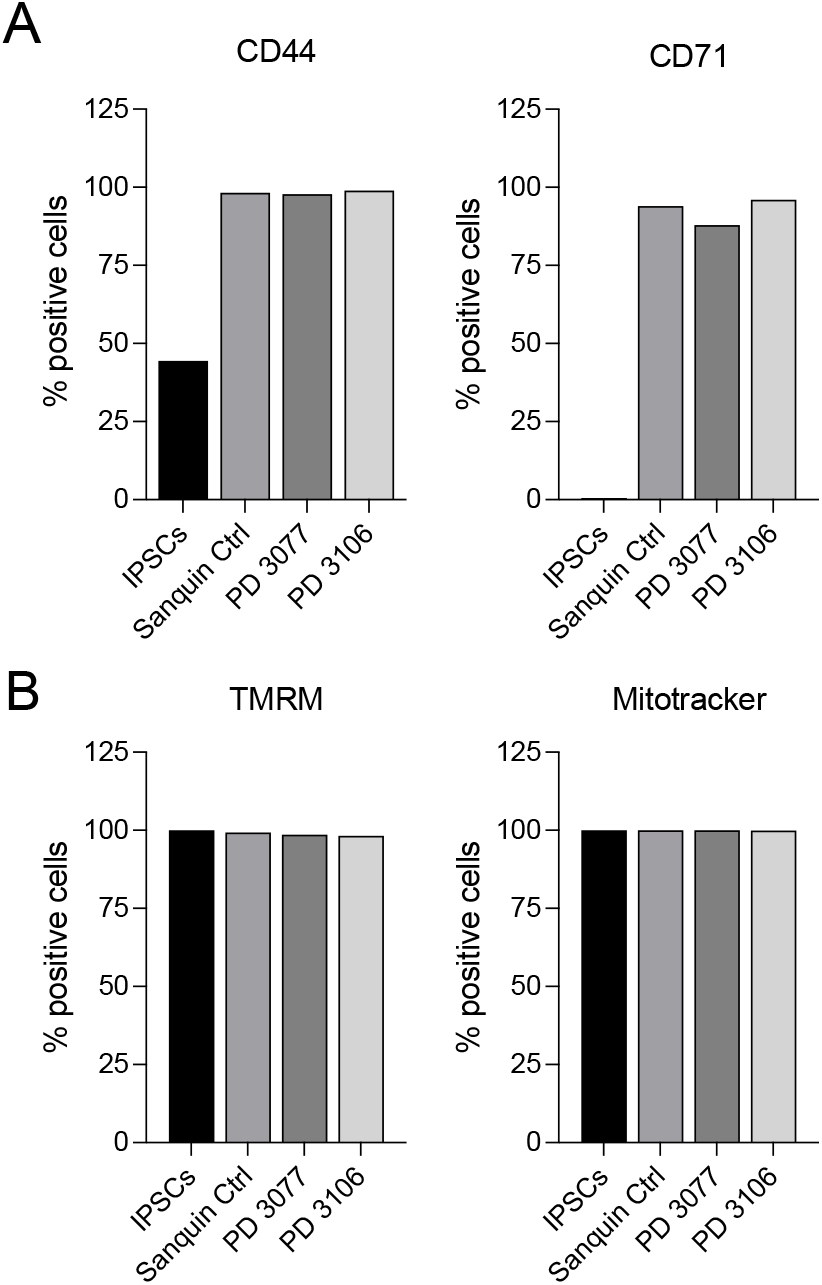
Cytofluorimetry analysis confirms the erythroblastic phenotype and the presence of functional mitochondria in cells from PD patients (PD3077, PD 3106) and healthy subjects. (**A**) As expected, isolated cells express the specific markers CD44, and CD71, and CD117. Negative control experiments in iPSCs fail to detect expression of the specific markers as expected. (**B**) In erythroblasts, mitochondria are polarized, as indicated by the TMRM signal.

We successfully isolated and expanded erythroblasts from blood human erythroid progenitors from PD patients and age matched donors (fig. 2). The erythroblast phenotype was confirmed by high level expression of the specific markers CD44 and CD71, which, as expected, displayed a significantly lower expression level in iPSc. MitoTracker and TMRM staining confirm that erythroblasts have functional, polarized mitochondria (fig. 2B).

We performed bioenergetic characterization of 35 erythroblasts lines of from PD patients with different disease severity (Milanese, Payan-Gomez et al. 2019) and from 18 lines from age and sex matched controls (Table 1).

Table of clinical severity

| SEVERITY 1 | SEVERITY 2 | SEVERITY 3 | SEVERITY 4 |
| --- | --- | --- | --- |
| **1032** | 1064 | 1042 | **152** |
| **3024** | 3007 | 3001 | 411 |
| 3043 | 3014 | 3013 | **1016** |
| 3054 | 3019 | 3020 | **1151** |
| 3098 | 3039 | 3038 | 2031 |
| 3122 | **3047** | **3085** | **3063** |
|  | 3074 | **3106** | 3097 |
|  | **3077** | **3115** | 3103 |
|  | 3086 | 3126 |  |
|  | 3120 | 3128 |  |
|  | 3160 |  |  |

As in our previous work, bioenergetic analyses was performed in standard glucose containing medium, i.e. in conditions allowing both glycolysis and oxidative phosphorylation, and in non-permissive conditions, where glucose was replaced by galactose. Because the use of galactose as a substrate in the glycolytic pathways is highly unfavored from the kinetic standpoint, the substrate cannot be metabolized through this pathway. Replacing glucose with galactose therefore forces bioenergetics through oxidative phosphorylation allowing detection of even subtle defects in mitochondrial function.

When we investigated bioenergetics in glucose containing medium, we detected significant reduction in both basal and maximal oxygen consumption (OCR) rates in PD erythroblasts (compare dark and light blue boxes in Fig 3). PD erythroblasts, however, can increase mitochondrial respiration if oxidative phosphorylation is the only available bioenergetic pathway, as it happens in galactose medium. Under these conditions, in fact, PD erythroblasts display higher basal respiration, maximal respiration, and respiration dedicated to ATP production than observed in glucose medium. (Fig. 3A). Conversely, increased mitochondrial activity in galactose medium was not observed in control erythroblasts (Fig. 3A). As expected, medium acidification (ECAR), which reflects glycolysis, is significantly reduced in galactose when compared to glucose medium in both healthy controls and PD patients (Fig 3B). These results reveal reduced respiration in PD specimens cultured in conditions that permit glycolysis. The comparison between glucose and galactose conditions, moreover, may discriminate between PD patients, which display different oxygen consumption rates in the two conditions, and healthy subjects.

**Figure 3.**
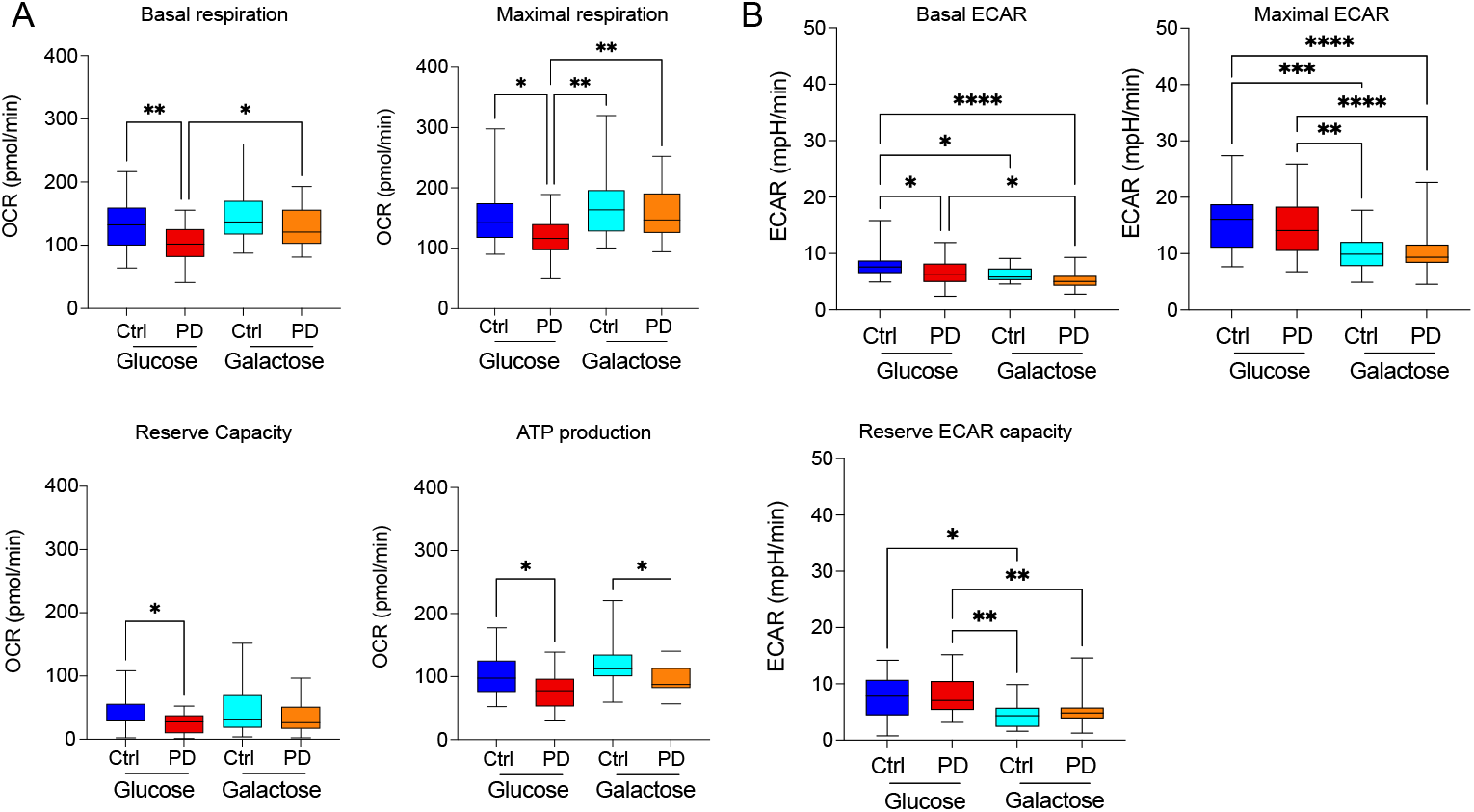
Bioenergetics profiles of erythroblasts isolated from iPD patients’ and age matched healthy controls. Galactose medium was used to force energetic metabolism through OXPHOS and to unmask mitochondrial defects. (**A**) In glucose conditions, PD patients display reduced basal and maximal oxygen consumption rate (compare dark and light blue boxes). Galactose medium increases OCR in PD erythronblasts (compare light blue and orange boxes), but not in cells from healthy controls (compare dark blue and red boxes). (**B**) ECAR is reduced In galactose medium.

In our previous studies in fibroblasts, we reported a correlation between the magnitude of bioenergetic defects and clinical severity, particularly with regards to non-dopaminergic symptoms (Milanese, Payan-Gomez et al. 2019). We therefore investigated whether biochemical parameters paralleled clinical measures also in erythroblasts originated from the very same patients we examined in our previous work. Patients were divided in four groups of increasing and comparable disease severity based on motor (UPDRS-III) and non-dopaminergic (SENS-PD) clinical scales, and upregulation of mitochondrial respiration in galactose conditions was higher in patients with more severe presentation (Milanese, Payan-Gomez et al. 2019) (suppl. Table 2).

When we compared mitochondrial bioenergetics among the four experimental groups, we found that maximal changed with disease severity more than basal respiration (fig.4). In glucose medium, maximal respiration was progressively reduced in the three groups with higher disease severity when compared to the control group, while it was unchanged patients with the mildest clinical presentation (fig. 4A) No differences were detected in basal respiration among the four groups (data not shown). Culturing cells in galactose medium caused an increase in maximal respiration, which reached statistical significance in the three groups with more severe presentation (fig. 4B). Higher values of the galactose/glucose OCR ratio in PD cells show, consistently with our previous data in fibroblasts (Milanese, Payan-Gomez et al. 2019), that respiration in PD cells can be increased in conditions forcing metabolism through OXPHOS and is therefore reversible. These elements support the concept that mitochondrial respiration suppression might reflect an adaptaptive, protective strategy to counteract pathogenic processes.

**Figure 4.**
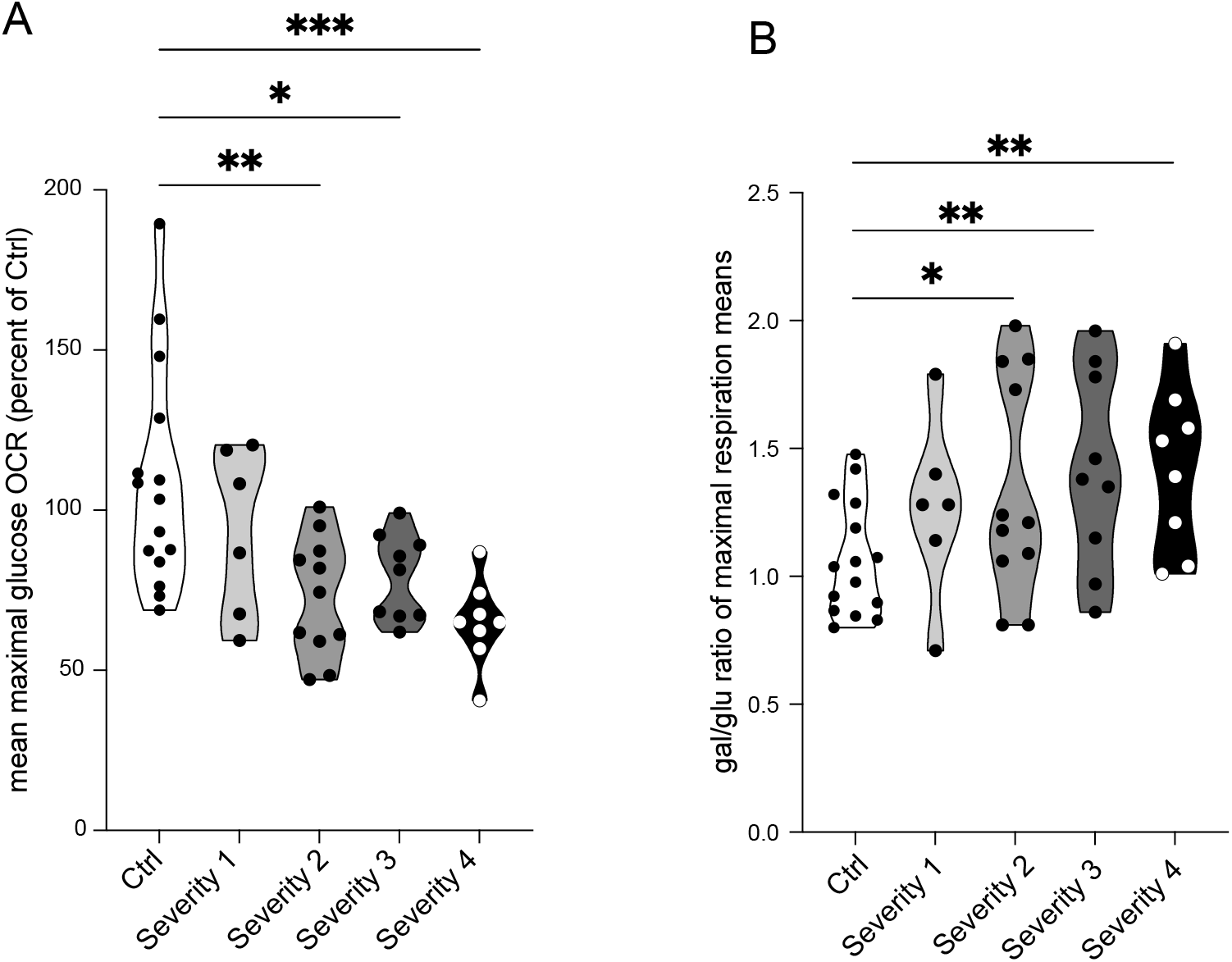
In erythroblasts, mitochondrial bioenergetics changes with the disease severity and the progression of the PD pathology. (**A**) Maximal respiration reveals differences among groups with different disease severity. Maximal respiration in glucose decreases with disease severity. (**B**) The ratio of OCR in galactose and glucose increases in PD and progressively augments with disease severity. One way ANOVA and Bonferroni post-hoc test.

Collectively, these results confirm that mitochondrial respiration is reduced in PD patients peripheral cells and that these alterations parallel disease severity

### iPSC derived dopaminergic neurons

We next extended our examination to dopaminergic neurons differentiated from inducible pluripotent stem cells derived (iDAN). We differentiated iDAN from iPS reprogrammed from fibroblasts from the same patients erythroblasts were originated from using small molecules (Grochowska, Carreras Mascaro et al. 2021). iPS expressed the expected phenotypic markers (fig. 5A). We analyzed iDAN from 16 idiopathic PD patients, thus a relatively large number of cell lines. As expected, iDAN expressed neuronal (MAP2) and dopaminergic (TH) markers (fig.5B).

**Figure 5.**
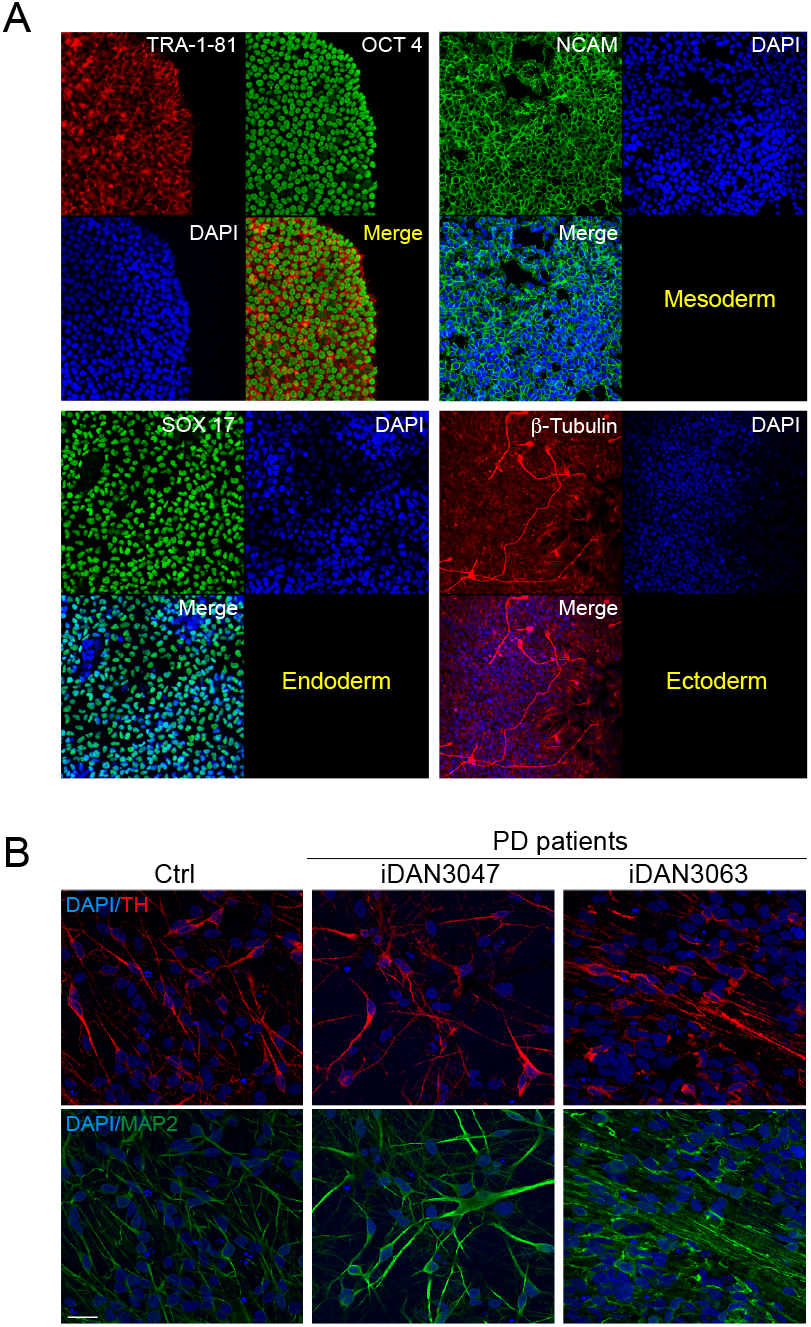
Fibroblasts from iPD patients’ and healthy controls have been reprogrammed into iPS (**A**) and differentiated into dopaminergic neurons (**B**). iPS cells have been characterized for self-renewability (**A**, left panel) and pluripotency (**A**, right panel). (**B**) Mature dopaminergic neurons have been checked for the expression of the tyrosine tydroxylase marker.

Bioenergetic analysis revealed significantly reduced basal mitochondrial respiration in PD lines when compared to healthy controls. Reserve capacity, however, was comparable between PD and healthy subjects (figure 6A). As expected, galactose medium remarkably reduced glycolysis, as indicated by reduced medium acidification (ECAR). Contrary to erythroblasts and fibroblasts, however, galactose medium did not lead to increased mitochondrial respiration. On the contrary, galactose medium induced a significant reduction in the reserve capacity levels in PD neurons (fig. 6A). As expected, ECAR was reduced in galactose medium (fig.6B).

**Figure 6.**
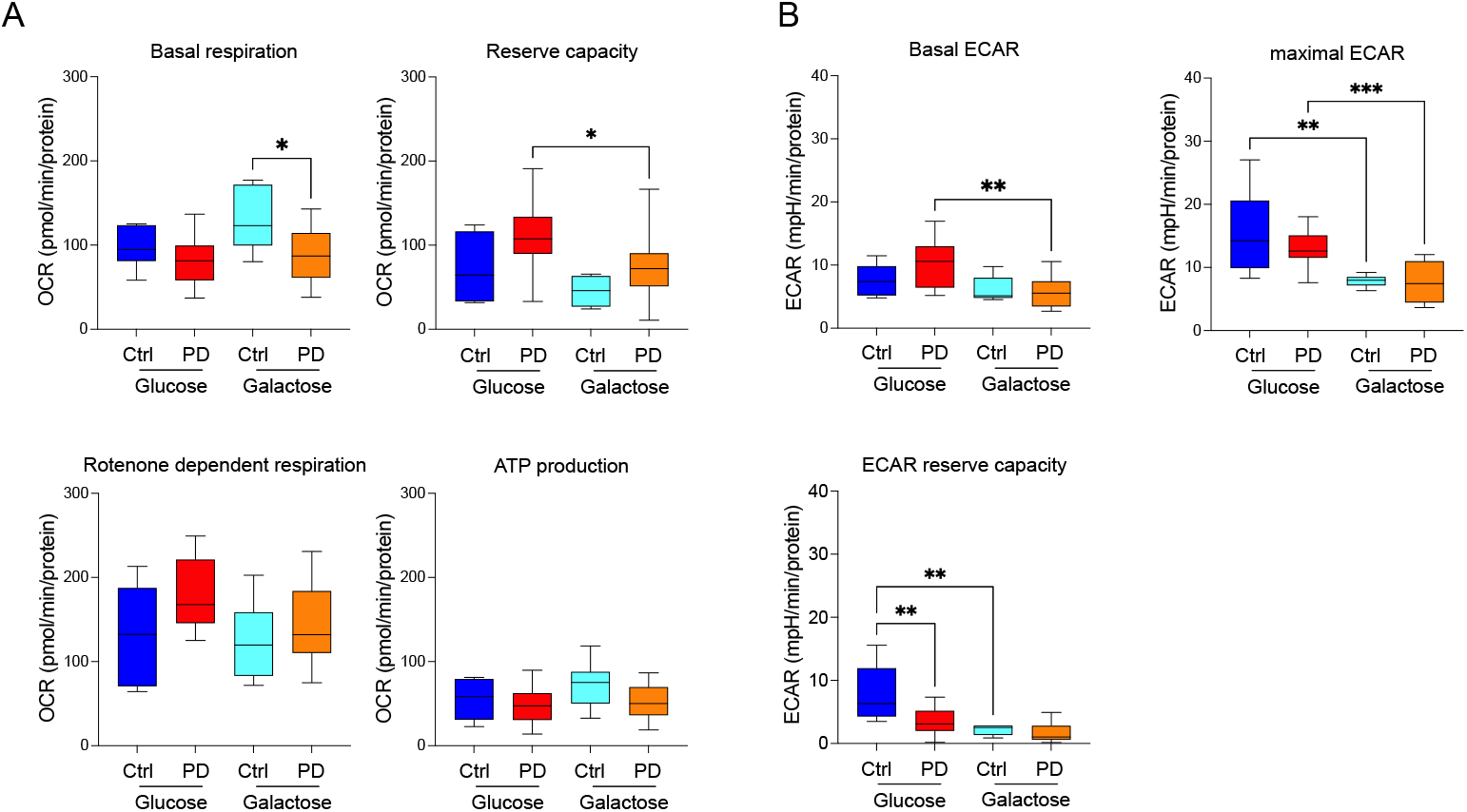
Bioenergetics profiles DA neurons derived from iPD fibroblasts patients’ and age matched healthy controls. Galactose medium was used to force energetic metabolism through OXPHOS and to unmask mitochondrial defects.

We found that also in iDAN maximal OCR in glucose increased with disease severity (fig. 7A). Galactose, however, was not able to increase respiration in iDAN (fig. 7B).

**Figure 7.**
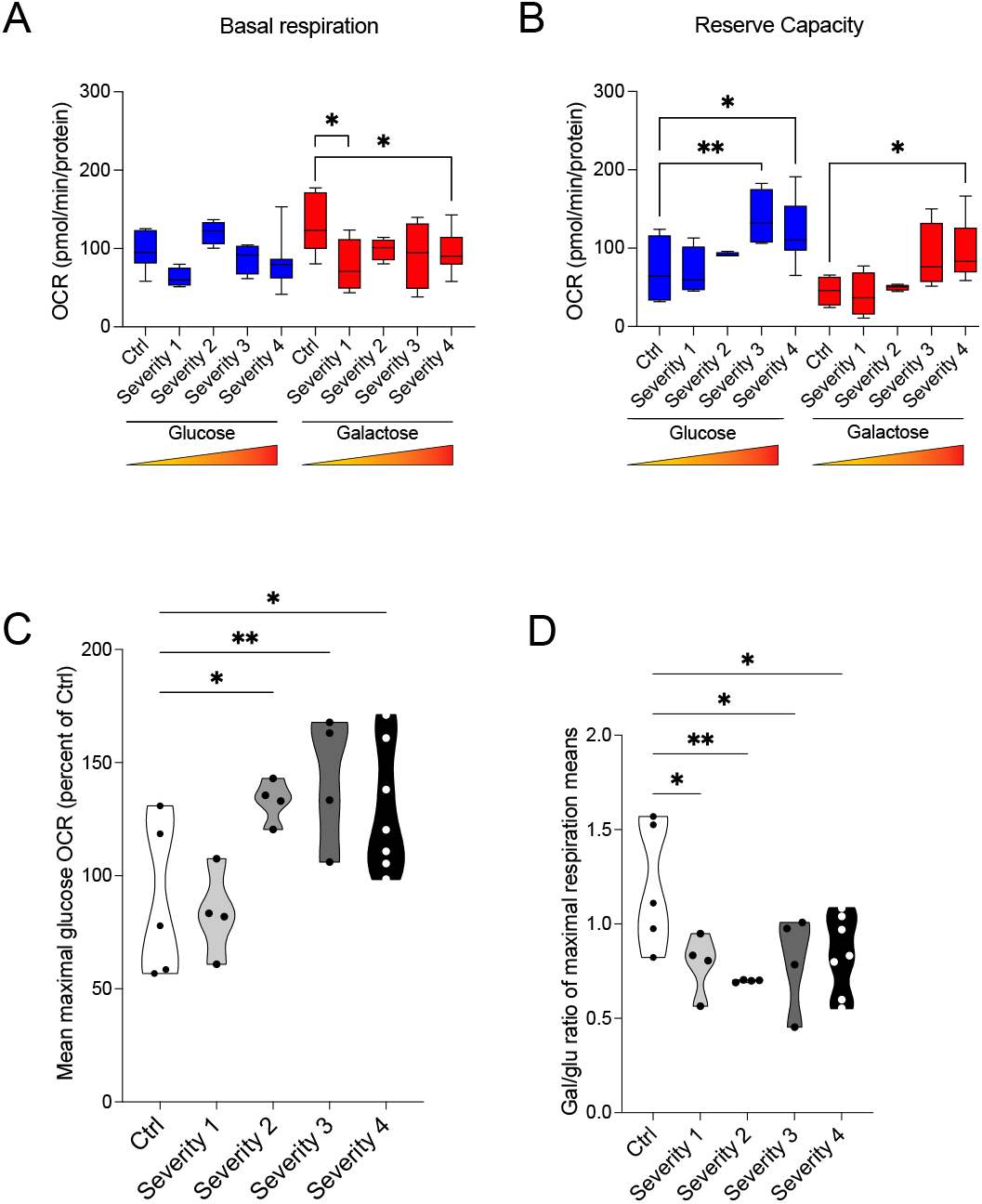
In DA neurons derived from iPD patients no direct correlation between mitochondrial bioenergetics disease severity and progression of the PD pathology is detected.

Collectively, these data show mitochondrial alterations in idiopathic PD at neuronal level and point to different bioenergetic features of cultured neurons versus peripheral cells such as erythroblasts and fibroblasts.

## Discussion

In this study, we present evidence confirming that bioenergetic defects in PD occur systemically, in peripheral cells, and are also detectable in dopamine neurons differentiated from iPS cells. PD erythroblasts display reduced mitochondrial respiration in culturing conditions allowing glycolysis, which can be however increased when cellular bioenergetics relies exclusively on mitochondrial respiration. These elements indicate that suppression of mitochondrial respiration in iPD is reversible and is consistent with previous findings in fibroblasts from our laboratory (Milanese, Payan-Gomez et al. 2019).

Mitochondrial biology in PD has been investigated in several cell types, including different PBMC subpopulations, fibroblasts, and human neurons (Mortiboys, Thomas et al. 2008, Mortiboys, Johansen et al. 2010, Seibler, Graziotto et al. 2011, Cooper, Seo et al. 2012, Zambon, Cherubini et al. 2019, Carling, Mortiboys et al. 2020). A strength of our study is that it analyzes and compares blood cells and iDAN derived from the same patient; additionally, it investigates an easily accessible blood cell type that to our best knowledge has been never studied in PD, i.e. erythroblasts. Moreover, studies in human neurons largely focused of PD familial forms so far, while we focused on idiopathic patients analyzing a rather large number of iDAN lines (N=17).

As expected, our data confirmed altered mitochondrial function in PD specimens and therefore substantiate the role of these organelles in PD pathobiology. The results also confirm that, at least in peripheral cell, alterations parallel clinical severity, consistently with data previously reported by us and other groups (Smith, Depp et al. 2018, Milanese, Payan-Gomez et al. 2019). Overall, our results corroborate the concept that peripheral cells may recapitulate neuronal pathology.

Blood samples are a highly convenient source of patient material given their accessibility and the minimally invasive procedures required to obtain the specimens. While several other studies analyzed the bioenergetic profile in PD blood cells, primarily in PBMC, to our best knowledge this is the first case that investigates mitochondrial function in erythroblasts. This approach offers some important conceptual and practical advantages. First, bioenergetics greatly differs among the various blood cells, and even among their subtypes. For instance, T-cell subtypes display large metabolic differences, being effector T-cells more reliant on glycolysis than regulatory and memory T-cells. The same applies to monocytes in their different polarized forms (O’Neill, Kishton et al. 2016). Moreover, erythroid cells express high level of a-syn – unlike other blood cells and fibroblasts - and may therefore be an amenable system to study the interplay between a-syn, bioenergetics, and other factors relevant for PD (Scherzer, Grass et al. 2008). Finally, erythroblasts can be prepared by amplifying in cell cultures a very small amount of starting material, generating a rather uniform population that does not require laborious and expensive cell sorting. Of note, other studies emphasized that the small number of obtainable blood cells limits the number of analyses that can be performed on a single sample (Smith, Depp et al. 2018); the use of expandable erythroblasts may obviate this issue, opening to further analytical options.

The evidence that PD erythroblasts can potentiate mitochondrial respiration when cultured in conditions that do not permit glycolysis, i.e. in galactose medium, replicates our previous findings in fibroblasts. Potentiation of mitochondrial respiration in galactose medium in PD peripheral cells suggests that in idiopathic PD, at least in a sub-group of cases, impaired mitochondrial function is reversible and may represent an adaptive protective response. Indeed, in our previous work we have shown that augmented mitochondrial respiration may favor alpha-synuclein stress (Milanese, Payan-Gomez et al. 2019). The possibility that suppression of mitochondrial respiration is a protective adaptation may be consistent with evidence obtained by other laboratories and by our group as well. An independent study has in fact shown a protective effect of the reversible inhibitor Mitochondrial Division Inhibitor 1 (Mdivi-1) in PD animal models (Bordt, Clerc et al. 2017). Moreover, we have shown that reversible and mild inhibition of complex I caused by its S-nitrosation is protective in multiple models of PD and improves bioenergetic efficiency in PD fibroblasts (Milanese, Payan-Gomez et al. 2019). A protective role for mitochondrial respiration suppression may also be consistent with recent hypotheses positing that in Alzheimer’s disease dysfunctional neurons increase mitochondrial respiration in a pathogenic mechanism that might be interpreted as an inverse-Warburg effect (Demetrius and Simon 2012, Demetrius, Magistretti et al. 2014). We deem the observation that reduction of mitochondrial respiration in PD particularly relevant because it might represent a paradigm shift from the general idea that this biochemical defect in PD is constitutive.

Differently than in peripheral cells, in iDNA potentiation of respiration in galactose medium was instead not detected. This finding might indicate that PD neurons may be more susceptible to mitochondrial defects, accordingly to a large body of literature showing that mild mitochondrial inhibition is particularly impactful on the dopaminergic system (Greenamyre, Cannon et al. 2010). It should be noted, however that iDAN cultures do not contain glial cells. Because neurons are metabolically coupled to astrocytes (Magistretti and Allaman 2015), this model is necessarily incomplete and additional experiments should be performed in co-cultures to determine whether this condition may permit potentiation of respiration when glycolysis is not permitted. Altogether, these elements reinforce the possibility that—at least in some subgroups of PD patients — mitochondrial function could be an amenable target for disease modification.

In conclusion, this offers additional arguments in favor of the use of peripheral cell in biomarker research for PD and confirms mitochondrial defects in human neurons derived from idiopathic patients.

## Acknowledgements

This study was funded by the Michael J. Fox Foundation (PGM)

## Contribution

Study design: PGM, SB, CM, MG. Experiments: SB, TL, LD, SF. Data analysis PGM, SB, CM, DS, CPG. PGM wrote the initial draft of the manuscript with input from all the authors. PGM conceived and directed the study.

